# Anterior cingulate cortex differently modulates fronto-parietal functional connectivity between resting-state and working memory tasks

**DOI:** 10.1101/722710

**Authors:** Xin Di, Heming Zhang, Bharat B Biswal

## Abstract

The brain fronto-parietal regions and the functional communications between them are critical in supporting working memory and other executive functions. The functional connectivity between fronto-parietal regions are modulated by working memory loads, and are shown to be modulated by a third brain region in resting-state. However, it is largely unknown that whether the third-region modulations remain the same during working memory tasks or were largely modulated by task demands. In the current study, we collected functional MRI (fMRI) data when the subjects were performing n-back tasks and in resting-state. We first used a block-designed localizer to define the fronto-parietal regions that showed higher activations in the 2-back than the 1-back condition. Next, we performed physiophysiological interaction (PPI) analysis using left and right middle frontal gyrus (MFG) and superior parietal lobule (SPL) regions, respectively, in three continuous-designed runs of resting-state, 1-back, and 2-back conditions. No regions showed consistent modulatory interactions with the seed pairs in the three conditions. Instead, the anterior cingulate cortex (ACC) showed different modulatory interactions with the right MFG and SPL among the three conditions. While increased activity of the ACC was associated with decreased functional coupling between the right MFG and SPL in resting-state, it was associated with increased functional coupling in the 2-back condition. The observed task modulations support the functional significance of the modulations of the ACC on fronto-parietal connectivity.

## 1. Introduction

Working memory involves distributed brain regions, most prominently the bilateral fronto-parietal network (Barch et al., 2013; Mencarelli et al., 2019; Owen, McMillan, Laird, & Bullmore, 2005). Understanding the functional integrations among the distributed regions is critical to understand the neural implementations of working memory. The bilateral fronto-parietal regions showed high correlations even in resting-state, thus forming lateralized fronto-parietal networks when using data driven methods such as independent component analysis (ICA) (Beckmann, DeLuca, Devlin, & Smith, 2005; Biswal et al., 2010; Di & Biswal, 2013). Because of the presence of functional connectivity during resting-state, it would be more critical to investigate the relative changes of functional connectivity during working memory tasks. Electroencephalogram (EEG) studies typically show increased connectivity in the theta band and reduced connectivity in the alpha band between fronto-parietal regions (Babiloni et al., 2004; Dai et al., 2017; Sauseng, Klimesch, Schabus, & Doppelmayr, 2005). As blood-oxygen-level dependent (BOLD) signals measured by functional MRI (fMRI), the signal synchronizations between some of the fronto-parietal regions were found to be reduced during higher working memory load condition compared with control condition, although these regions were more activated in the same contrast (Di & Biswal, 2019).

In addition to task modulations, functional connectivity between two regions might also be modulated by a third region (Di & Biswal, 2015a; Friston et al., 1997). In the context of working memory, some executive or distractive signals from other brain region might facilitate or disrupt the functional communications between fronto-parietal regions. This will result in higher order interactions among three brain regions, which can be studied using physiophysiological interaction (PPI) model (Di & Biswal, 2013; Friston et al., 1997) or nonlinear dynamic causal modeling (Stephan et al., 2008). Several studies have been performed to characterize the modulatory interactions in resting-state (Di & Biswal, 2013, 2014, 2015a, 2015b). Particularly, we defined the fronto-parietal regions of interest (ROIs) by using ICA and performed PPI analysis on the left and right fronto-parietal ROIs, respectively (Di & Biswal, 2013). We identified several medial frontal and parietal regions that showed negative modulatory interaction with the fronto-parietal ROIs, indicating that the increases of activity of these regions are accompanied by reduced fronto-parietal functional connectivity. However, this analysis was only performed in resting-state. It remains unclear whether similar effects would be shown in task conditions, or it could alter significantly upon task demands.

The goal of the current study is to examine whether modulatory interactions of the fronto-parietal regions are modulated by task demands. We adopted a n-back paradigm with varying working memory loads where the bilateral fronto-parietal regions are consistently activated (Barch et al., 2013; Owen et al., 2005). We first used a block-designed localizer to identify the fronto-parietal regions that showed higher activations during the 2-back than the 1-back condition. We then performed PPI analysis by using the frontal and parietal ROIs in three separate continuous task conditions, i.e. resting-state, 1-back, and 2-back conditions. We examined two competing hypotheses. First, there are modulatory interactions of a third region with the two ROIs, and the effects are consistent across the conditions. In contrast, there may be modulatory interactions of a third region with the two ROIs, but the effects highly depend on the task conditions. We performed conjunction analysis to identify brain regions that may fulfill the first hypothesis, and performed repeated measure one-way ANOVA to find regions that may fulfill the second hypothesis.

## 2. Methods

### 2.1. Subjects

Fifty participants (26 females) were recruited for the current study. The mean age was 22.34 years (19 – 24 years, SD = 1.303). One subject was removed because of large head motion during MRI scan. All participants reported normal auditory and normal or corrected-to-normal visual acuity, and were free of neurological or psychiatric problems. All study procedures were carried out with written informed consent of each subject. Each subject received honorarium of 200 RMB for the participation. The study was approved by institutional review board.

### 2.2. Study procedure

At the beginning of the MRI scan session, the participants underwent a resting-state fMRI scan (8 min 30 sec). The participants were instructed to lay still with eyes open and staring at a white cross fixation on a dark background. Four working memory task runs were then performed with the following order: two block-designed runs with both 1-back and 2-back condition in each run (3 min 46 sec each), one continuous run of 1-back condition (5 min 10 sec), and one continuous run of 2-back condition (5 min 10 sec). A high resolution anatomical T1-weighted MRI was scanned at the end of the MRI session.

#### 2.2.1. N-back task

The n-back task tests the participants’ working memory on the spatial locations of letters presented on the screen. A white cross fixation was presented at the center of the dark screen throughout the experiment. A random letter would be presented in 1 of the 4 visual field quadrants around the fixation. In a n-back task condition (n =1 or 2), participants were asked to press the left button with the left thumb when the location of the current letter matched with the one presented “n” item(s) back, and pressed the right button with the right thumb when it didn’t match with the one presented “n” item(s) back. The letter stimulus was presented for 500 ms, followed by an interstimulus interval of 2500 ms. One third of the total trials were “matches”. Participants were instructed to focus only on the location of the letter, but not on the letter itself, and to classify the stimuli as accurately and quickly as possible. Visual stimuli were presented and responses were collected using E-Prime (Psychology Software Tools).

The n-back task procedures were designed in two ways. First, in the two localizer runs, the n-back stimuli were presented as separate blocks of 1-back or 2-back conditions. Each run started with a 10 s fixation. Then, each of the block consisted of 8 trials (24 sec), with a 24-s fixation period intercepted between the task blocks. The orders of task blocks of the two runs were “ABBA” and “BAAB”, respectively. As a result, each run lasted for 3 min and 46 sec. Second, in the two continuous runs, the n-back trials were presented continuously without long fixation period between them. The 1-back and 2-back conditions were allocated in two separate runs. Each run started with a 10 s fixation period, followed by 100 trials. Each run lasted for 5 min and 10 sec.

#### 2.2.2. MRI scanning parameters

MRI data were acquired on a 3T GE Signa Scanner (General Electric Company, Milwawkee, WI) in functional MRI center at University of Electronic Science and Technology of China. An 8-channel head coil was used. The scanning parameters for the fMRI were: TR (repetition time) = 2000 ms; TE (echo time)) = 30 ms; flip angle = 90°; FOV (field of view) = 240×240 mm^2^; matrix size = 64×64; axial slice number = 42 with slice thickness = 3 mm and gap = 0). As a result, each resting-state run was consisted of 255 images, each block-designed run was consisted of 113 images, and each continuous task run was consisted of 155 images. Structural T1-weighted images were acquired using the following parameters: TR = 6 ms; TE = Minimum; TI = 450 ms; flip angle = 12°; FOV = 256×256 mm^2^; matrix size = 256×256; sagittal slice number = 156 with slice thickness = 1 mm.

### 2.3. FMRI data analysis

#### 2.3.1. Preprocessing

FMRI images were processed using SPM12 (SPM, RRID: SCR_007037; https://www.fil.ion.ucl.ac.uk/spm/) under MATLAB environment (R2017b). The anatomical image of each subject was segmented into gray matter (GM), white matter (WM), cerebrospinal fluid (CSF), and other brain tissue types, and normalized into standard Montreal Neurological Institute (MNI) space. The first five functional images of each run were discarded from analysis. The remaining images were realigned to the first image of each run, and coregistered to the anatomical image. The deformation field images obtained from the segmentation step were used to normalize all the functional images into MNI space, with a resampled voxel size of 3 × 3 × 3 mm^3^. All the images were spatially smoothed using an 8 × 8 × 8 mm^3^ Gaussian kernel.

We calculated frame-wise displacement for the translation and rotation directions to reflect the amount of head motions (Di & Biswal, 2015a). We adopted the threshold of maximum frame-wise displacement of 1.5 mm or 1.5 degree (half voxel size), or mean frame-wise displacement of 0.2 mm or 0.2 degree. The subjects with any of the five runs exceeding the threshold would be removed from the analysis. As a result, one subject’s data were discarded.

#### 2.3.2. Activation analysis of the block-designed runs

We first defined general linear model (GLM) to perform voxel-wise analysis on the block-designed runs to identify task activations between the 2-back and 1-back conditions. The two runs were modeled together with their own task regressors, covariates, and constant terms. The 2-back and 1-back conditions were defined as two box-car functions convolved with canonical hemodynamic response function (HRF). The first eigenvector of the signals in the WM and CSF, respectively, and 24 head motion regressors (Friston, Williams, Howard, Frackowiak, & Turner, 1996) were added as covariates. There was also a high-pass filter (1/128 Hz) implicitly implemented in the GLM. After model estimation, a contrast of 2-back – 1-back was defined to reflect the differences of activations between the two conditions.

Group level analysis was performed using one sample t test GLM with the contrast images of 2-back vs. 1-back as dependent variables. Activated clusters were first identified using a threshold of p < 0.001 of two-tailed t test (Chen et al., 2019), and the cluster extent was thresholded at cluster level false discovery rate (FDR) of p < 0.05. Because we were interested in fronto-parietal regions, we searched the peak coordinates of the resulting clusters as well as local maxima within large clusters that covered these regions. As a result, we defined bilateral middle frontal gyrus regions (MNI coordinates: RMFG, 24, 11, 56; LMFG, −24, 8, 50) and superior parietal lobule (MNI coordinates: LSPL, −18, −70, 50; RSPL, 21, −67, 53) as ROIs.

#### 2.3.3. Physiophysiological interaction analysis of the continuous-designed runs

We first defined GLMs for each continuous run and subject to define ROIs. The GLMs only included the WM/CSF, head motion, and constant regressors, but did not include any task regressors. A high-pass filter (1/128 Hz) was also implicitly implemented in the GLM. After model estimation, the time series of the LMFG, LSPL, RMFG, and RSPL were extracted within spherical ROIs of 6 mm radius centered at the above mentioned MNI coordinates. All the effects of no-interests, i.e. WM/CSF signals, head motion, constant, and low-frequency drifts were adjusted during the time series extraction. PPI terms were calculated for LMFG and LSPL, and RMFG and RSPL, respectively. The time series of the two ROIs were deconvolved with canonical HRF, multiplied together, and convolved back with HRF to form a PPI term (Di & Biswal, 2013; Gitelman, Penny, Ashburner, & Friston, 2003). Here we only focused on within hemisphere fronto-parietal connectivity, e.g. LMFG and LSPL, but excluded inter-hemisphere connectivity, e.g. LMFG and RSPL. This is because usually there is no direct anatomical connection between two different regions across hemispheres. The observed functional interactions between them, e.g. LMFG and RSPL, are usually mediated by one of their corresponding region in the opposite hemisphere, e.g. RMFG or LSPL.

Next, new GLMs were built with the time series of the two ROIs and the PPI term between them for each of the ROI pairs and task conditions. Other regressors of no-interests as well as the implicit high-pass filter were also included in the GLMs. The beta estimates corresponding to the interaction term was the effect of interest, which were used for the group level analysis. We note that the beta estimates are not a function of sample size (the number of time points in this case). Therefore, the comparisons of betas between resting-state and n-back runs are not biased by the differences in time points.

The first goal of the group analysis is to identify regions that show modulatory interaction effects consistently present in the three conditions. We performed conjunction analysis of the three conditions. First, second-level GLMs were built for the LMFG-LSPL and RMFG-RSPL analyses, respectively, using a one-way analysis of variance (ANOVA) model implemented in SPM. The GLM included three columns representing the three conditions. Second, a t contrast was defined for each condition for the positive and negative directions, respectively. Finally, we examined the conjunction effects of the three conditions for the positive and negative effects, respectively, using a threshold of one-tailed p < 0.0005 (corresponding to two-tailed p < 0.001). Cluster level FDR of p < 0.05 was used for the cluster extent threshold. Because there were no clusters survived at the two-tailed p < 0.001 threshold, we also explored lower threshold of two-tailed p < 0.01 for potential effects.

The second goal is to identify regions that showed variable modulatory interactions in the three conditions. Repeated measure one-way ANOVA model was used for this purpose, with the three conditions as three levels of a factor. The significant results of the repeated measure ANOVA indicate differences in the PPI effects between any two of the three conditions. The resulting statistical maps were thresholded at p < 0.001 with cluster level FDR at p < 0.05.

## 3. Results

### 3.1. Task activations in the localizer runs

We observed typical bilateral fronto-parietal regions that showed higher activations during the 2-back condition compared with 1-back condition (Figure 1 and Table 1). The frontal clusters mainly covered the bilateral middle frontal gyrus and precentral gyrus. The parietal clusters mainly covered the bilateral superior parietal lobule and precuneus. The right cerebellum and left basal ganglia were also activated. There were also reduced activations in the 2-back compared with 1-back condition, mainly in the default model network and bilateral temporo-opercular regions.

**Table 1.** Clusters that showed increased or decreased activations in the 2-back condition compared with the 1-back condition in the block designed runs. The cluster was defined as two tailed p < 0.001, with cluster level false discovery rate (FDR) of p < 0.05.

| p (cluster FDR) | voxels | Coordinates |  |  | peak T | Label |
| --- | --- | --- | --- | --- | --- | --- |
|  |  | x | y | z |  |  |
| < 0.001 | 2108 | 24 | 11 | 56 | 11.65 | Right middle frontal gyrus |
|  |  | -24 | 8 | 50 | 10.72 | Left middle frontal gyrus |
|  |  | -48 | 5 | 32 | 9.810 | Left precentral gyrus |
| < 0.001 | 2897 | -6 | -61 | 44 | 10.73 | Precuneus |
|  |  | -18 | -70 | 50 | 10.68 | Left superior parietal lobule |
|  |  | 21 | -67 | 53 | 10.44 | Right superior parietal lobule |
| 0.004 | 120 | 48 | 5 | 23 | 7.00 | Right precentral gyrus |
| 0.003 | 149 | 27 | -61 | -37 | 6.92 | Right cerebellum |
|  |  | 9 | -73 | -31 | 4.78 | Right cerebellum |
| 0.003 | 136 | -18 | 5 | 11 | 5.84 | Left caudate |
|  |  | -30 | 26 | 2 | 5.75 | Left anterior insula |
| 0.038 | 63 | -33 | 50 | 2 | 4.20 | Left middle frontal gyrus |
|  |  | -42 | 50 | 2 | 4.02 | Left middle frontal gyrus |
| < 0.001 | 661 | -3 | -16 | 32 | -8.73 | Middle cingulate gyrus |
|  |  | 0 | -37 | 20 | -6.08 | Posterior cingulate gyrus |
|  |  | 0 | -28 | 44 | -5.42 | Posterior cingulate gyrus |
| < 0.001 | 660 | 39 | -19 | 20 | -6.54 | Right parietal operculum |
|  |  | 36 | -16 | 2 | -5.56 | Right posterior insula |
|  |  | 39 | 2 | -1 | -5.16 | Right anterior insula |
| < 0.001 | 910 | 12 | 59 | 20 | -6.11 | Superior frontal gyrus |
|  |  | -6 | 62 | 8 | -5.86 | Medial superior frontal gyrus |
|  |  | -9 | 53 | -1 | -5.84 | Medial superior frontal gyrus |
| < 0.001 | 498 | -36 | -10 | -4 | -5.39 | Left posterior insula |
|  |  | -63 | -25 | 5 | -4.73 | Left superior temporal gyrus |
|  |  | -39 | -19 | 17 | -4.64 | Left central operculum |
| 0.037 | 74 | 21 | 38 | -1 | -5.19 | Anterior cingulate gyrus |

X, y, and z coordinates are in (Montreal Neurological Institute) MNI space.

**Figure 1.**
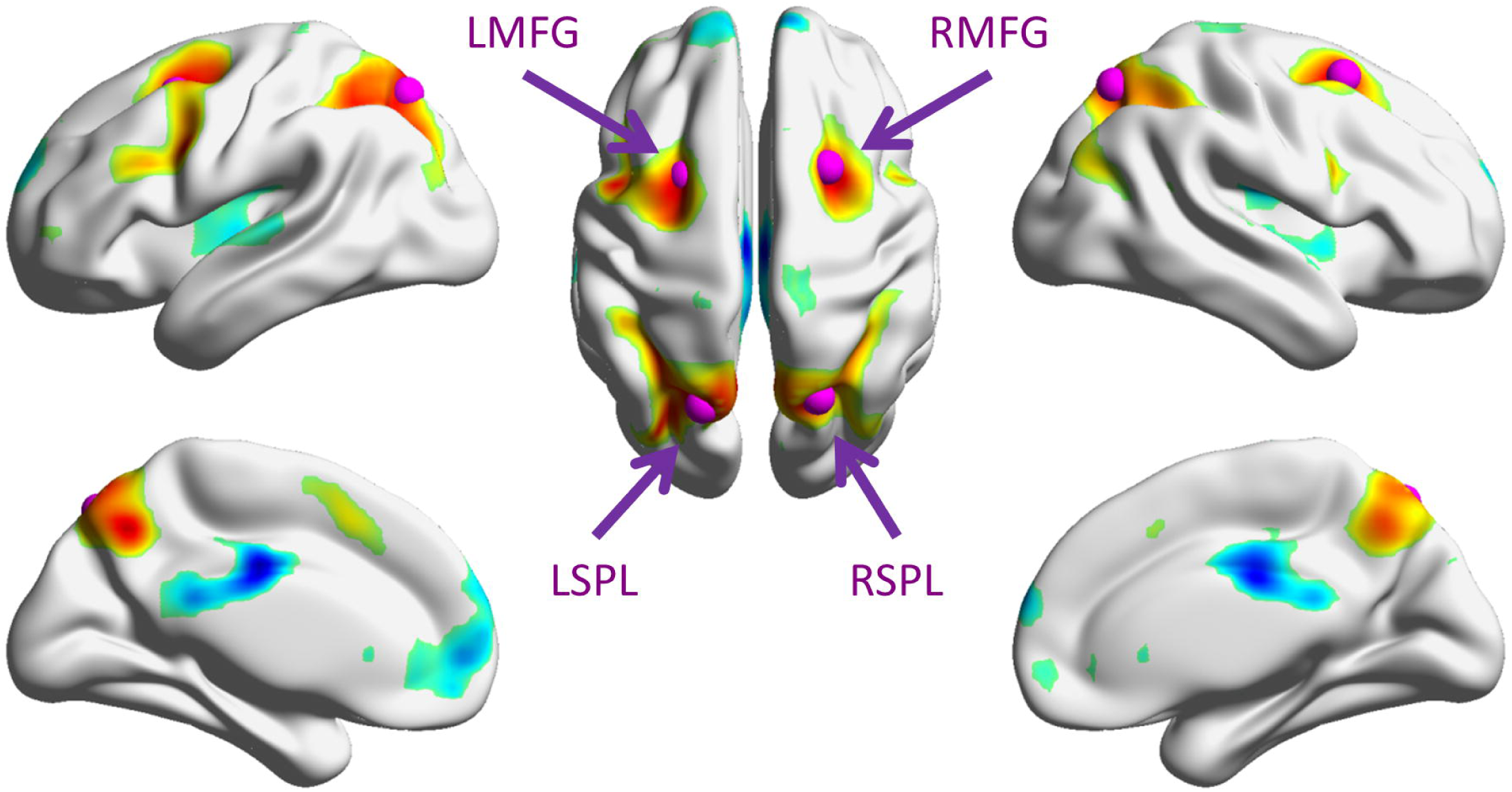
Increased (warm color) and decreased (cold color) activations in the 2-back condition compared with the 1-back condition. The map was thresholded at p < 0.001 (two-tailed) with cluster-level false discovery rate of p < 0.05. The purple spheres illustrate the four regions of interest used in the physiophysiological interaction (PPI) analysis. The surface presentation was made by using BrainNet Viewer (RRID: SCR_009446) (Xia, Wang, & He, 2013). LMFG, left middle frontal gyrus; RMFG, right middle frontal gyrus; LSPL, left superior parietal lobule; and RSPL, right superior parietal lobule.

### 3.2. Modulatory interactions during different task conditions

We first performed conjunction analysis to identify regions that showed consistent PPI effects across the three conditions. No statistical significant clusters were found of any sizes at p < 0.001 for both the LMFG-LSPL and RMFG-RSPL analyses. We further checked the threshold of p < 0.01, and still there were no clusters of any sizes survived.

Repeated measure one-way ANOVA showed only significant effects on the modulatory interactions of RMFG and RSPL. As shown in Figure 2 and Table 2, the only cluster mainly covered the anterior cingulate cortex (ACC). The cluster-level FDR corrected p value (0.005) also survived Bonferroni correction for the two analyses (RMFG/RSPL and LMFG/LSPL). Post-hoc analysis showed that the PPI effect in the ACC was positive in the 2-back condition but negative during resting-state (Figure 2B). And the differences among the three conditions were mainly driven by the differences between the 2-back condition and the other two conditions. Repeated measure one-way ANOVA of the modulatory interactions of LMFG and LSPL showed a similar cluster in the ACC. However, the cluster size could not pass the cluster-level threshold.

**Table 2.** Cluster that showed different physiophysiological interaction (PPI) effects with right middle frontal gyrus (RMFG) and right superior parietal lobule (RSPL) among the resting-state, 2-back, and 1-back conditions in the continuous runs (repeated measure one way analysis of variance, ANOVA). The cluster was defined as p < 0.001, with cluster level false discovery rate (FDR) of p < 0.05.

| p (cluster FDR) | voxels | Coordinates |  |  | peak F | Label |
| --- | --- | --- | --- | --- | --- | --- |
|  |  | x | y | z |  |  |
| 0.005 | 133 | -3 | 32 | 14 | 14.94 | Anterior cingulate gyrus |
|  |  | 9 | 35 | 5 | 14.82 | Anterior cingulate gyrus |
|  |  | 3 | 44 | -4 | 8.27 | Anterior cingulate gyrus |

X, y, and z coordinates are in (Montreal Neurological Institute) MNI space.

**Figure 2.**
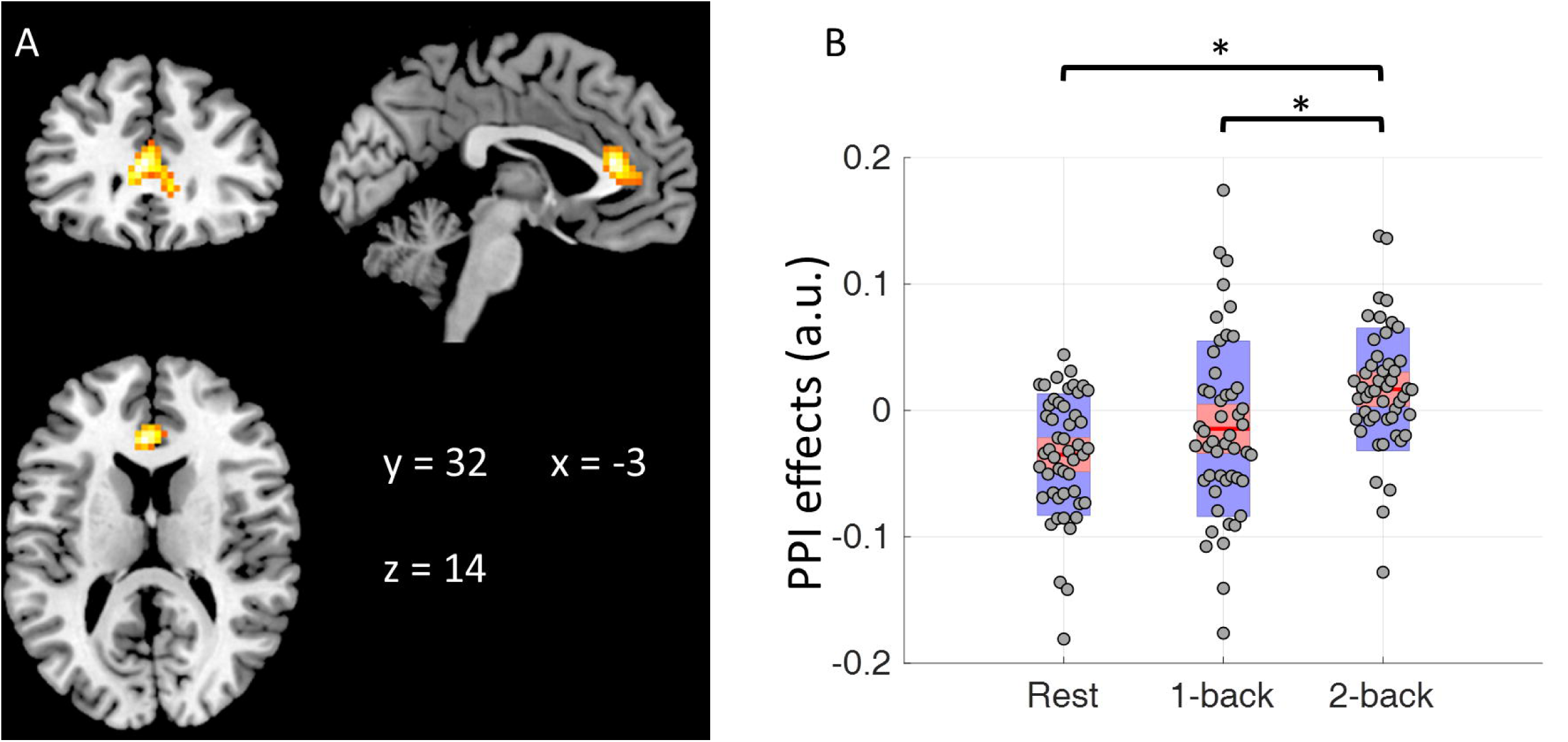
A) Region that showed different modulatory interactions with right middle frontal gyrus (RMFG) and right superior parietal lobule (RSPL) among the three task conditions (repeated measure one way analysis of variance, ANOVA). The map was thresholded at p < 0.001 with cluster level false discovery rate (FDR) of p < 0.05. B) Mean modulatory interactions of the cluster in the in the three conditions. The center red lines represent the mean effects, and the light red bars and light blue bars represent 95% confidence interval and standard deviation, respectively. * indicates statistical significance in post-hoc pair-wise comparisons at p < 0.05. Panel B was made by using notBoxPlot (https://github.com/raacampbell/notBoxPlot). A.u., arbitrary unit.

In order to better interpret the PPI effects in the ACC, we correlated the mean PPI effects in the ACC cluster with RMFC and RSPL with behavioral measures of mean reaction time and accuracy (Figure 3). The PPI effect showed a very small correlation with reaction time (*r = −0.16*), and a moderate negative correlation with the accuracy (*r = −0.39*). But it can be seen in Figure 3C that there were potential outliers near the x axis that might introduce spurious correlations. We therefore performed bootstrapping for 10,000 times to obtain a 95% confidence interval of the correlation (−0.6352, 0.0046) (Figure 3D).

**Figure 3.**
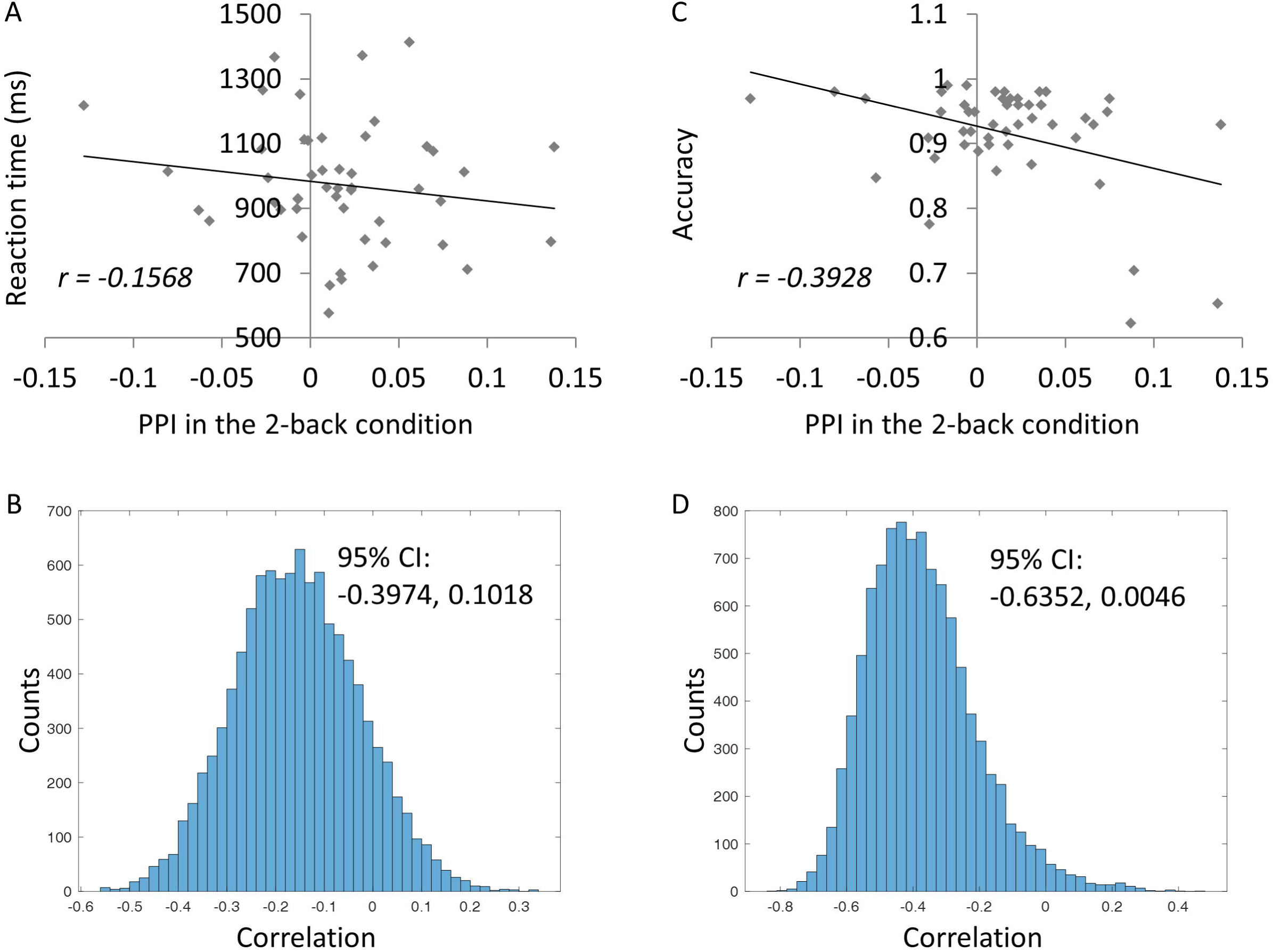
Behavioral correlates of the mean modulatory interactions in the anterior cingulate cortex (ACC) with right middle frontal gyrus (RMFG) and right superior parietal lobule (RSPL) during the 2-back continuous run. A and B illustrate the relations between the modulatory interactions and reaction times and 10,000 bootstrapping distributions of the correlations. C and D illustrate the relations between the modulatory interactions and accuracy and 10,000 bootstrapping distributions of the correlations.

### 3.3. Post hoc task activation analysis

Lastly, we also extracted the mean task activations of the ACC in the block-designed runs (Figure 4). The ACC showed reduced activations in both the 1-back and 2-back conditions with reference to the fixation baseline. But the activations were more negative in the 2-back condition than in the 1-back condition (paired t test: *t(48) = 4.49, p < 0.001*).

**Figure 4.**
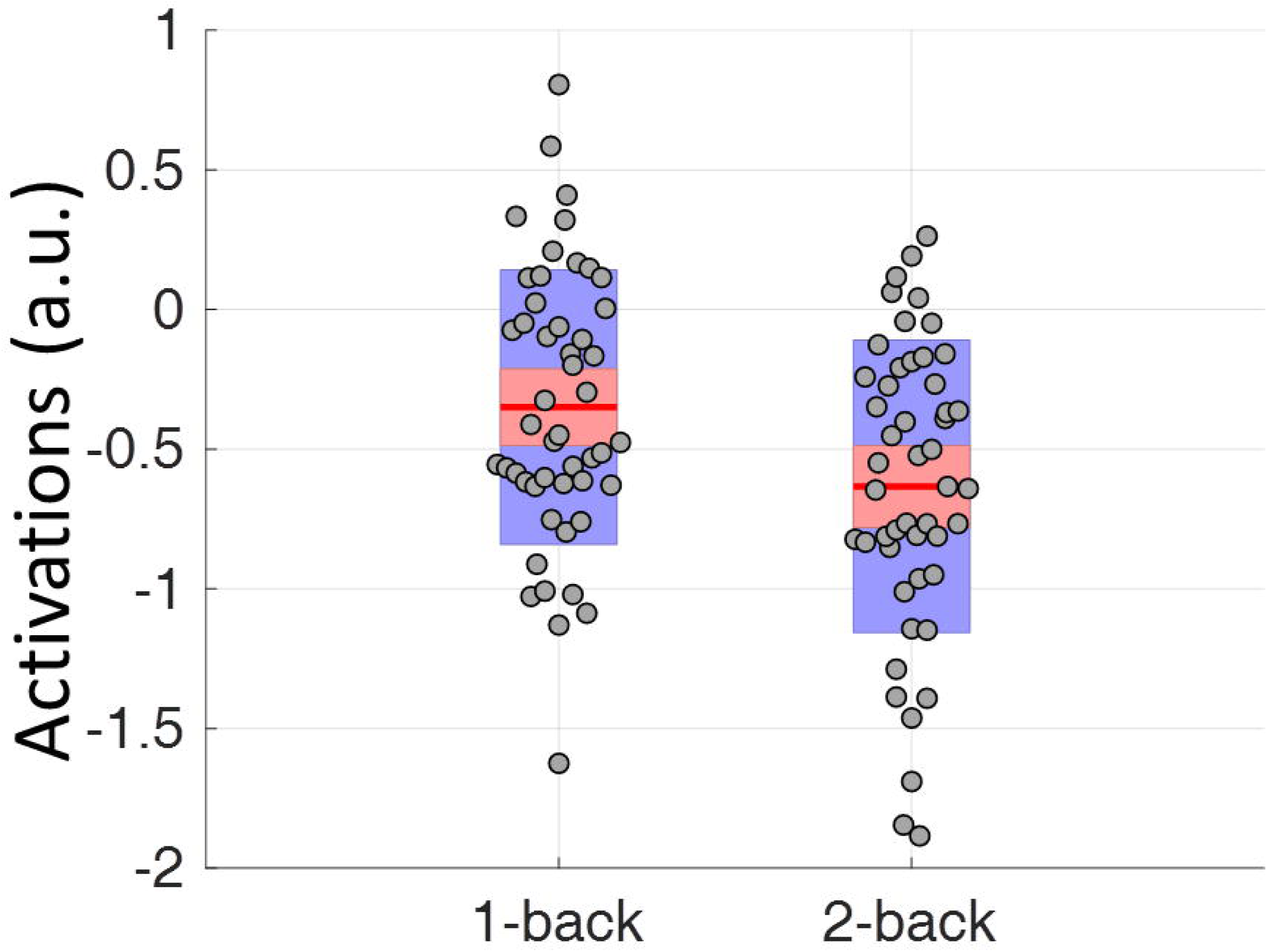
Mean task activations of the anterior cingulate cortex (ACC) cluster in the block-designed runs. The center red lines represent the mean effects, and the light red bars and light blue bars represent 95% confidence interval and standard deviation, respectively. This figure was made by using notBoxPlot (https://github.com/raacampbell/notBoxPlot). A.u., arbitrary unit.

## 4. Discussion

By comparing modulatory interactions of two key regions in working memory across three continuously designed task conditions, the current analysis identified the ACC that showed different modulatory interactions with the RMFG and RSPL in the resting-state, 1-back, and 2-back conditions. On the other hand, no regions showed consistent modulatory interactions with the fronto-parietal regions across the three conditions. The activity in the ACC was positively correlated with the connectivity of RMFG and RSPL during the 2-back condition, but was negatively correlated with the connectivity of RMFG and RSPL in resting-state. Due to the nature of regression model, this is impossible to infer the directions of the modulations (Di & Biswal, 2013). However, the RMFG and RSPL were co-activated by the working memory task and are also considered part of the same functional network (Biswal et al., 2010; Yeo et al., 2011), while the ACC was more deactivated in the 2-back condition. We therefore prefer to interpret the results as that the ACC increase the functional connectivity between RMFG and RSPL during the 2-back condition, and reduce the functional connectivity between the RMFG and RSPL.

Due to the fact that the ACC was negatively activated in the task conditions compared with the fixation condition (Figure 4), it is likely that the ACC is part of the default mode network (Raichle et al., 2001). The current PPI results are consistent with our previous study in resting-state, which also showed some midline regions from the default mode network having negative modulatory interactions with RMFG and RSPL (Di & Biswal, 2013). The task positive network including the fronto-parietal regions and the default mode network are anti-correlated both in resting-state (Fox et al., 2005) and during task executions (Shulman et al., 1997). The current results together with our previous work (Di & Biswal, 2013) further confirm that the competing nature of the task positive and default mode networks not only exist in first order relationships but also in higher order interactions.

More interestingly, current analysis found that the modulatory interactions among ACC, RMFG, and RSPL were largely modulated by task conditions. In contrast to the resting-state, the ACC showed no significant modulatory interactions in the 1-back condition, and positive modulatory interactions in the 2-back condition. The task dependent effect is in line with some studies that have demonstrated task modulated modulatory interactions in other brain systems by using higher order psycho-physio-physiological interaction models (Gorka, Knodt, & Hariri, 2015; Stamatakis, Marslen-Wilson, Tyler, & Fletcher, 2005). In neuronal level models, it has also been shown that higher order interactions present only in certain task conditions (Ganmor, Segev, & Schneidman, 2011; Macke, Opper, & Bethge, 2011). Taken together, all the evidence conversely suggests that high order interactions may be sensitive to task demands.

During the 2-back condition with higher working memory loads, the signals from the ACC were associated with increased functional communications between the fronto-parietal regions. One of the functions of the ACC is error detection and conflict monitoring (Bush, Luu, & Posner, 2000). Then, the ACC activity may represent error related signals that would enhance the communications between the fronto-parietal regions to maintain task performances. The brain-behavioral correlation analysis supported this interpretation. The modulatory interactions in the 2-back condition were not correlated with reaction time, but were negatively correlated with accuracy. In other words, the more errors one made, the larger the modulatory interactions were among ACC, RMFG, and RSPL.

The current study adopted functionally defined ROIs of the MFG and SPL from a localizer for the PPI analysis. The bilateral MFGs are a little anterior to the premotor regions and posterior to the dorsolateral prefrontal cortex reported in a meta-analysis of n-back tasks (Owen et al., 2005). And the bilateral SPLs are superior and posterior to the inferior parietal lobule region reported in (Owen et al., 2005). The differences may represent discrepancies in task designs and control conditions. But the fact that these regions showed the highest contrast between the 2-back and 1-back condition in the current localizer task support the usage of these regions to represent regions that are involved in working memory process. The fronto-parietal ROIs also do not exactly match with those used in the resting-state study (Di & Biswal, 2013). But similar to this paper, the current analysis showed negative modulatory interactions in the middle line region of ACC with RMFG and RSPL (Di & Biswal, 2013).

The current analysis adopted a ROI-based approach, with ROIs identified directly from the working memory task studied. This helped us to focus on specific brain regions that are related to the task. The whole brain PPI analysis identified a region that are not a part of the fronto-parietal network nor activated during the working memory tasks. It is reasonable because our previous study has shown that modulatory interactions are more likely to take place among regions from different brain networks (Di & Biswal, 2015a). There may be other brain regions that involve higher order interactions with one of the fronto-parietal regions. But the potential interactions will increase exponentially when considering the combinations of two brain regions outside the fronto-parietal network, making it difficult to do an exhaustive search based on the current sample size. Further studies may adopt the whole brain approach (Di & Biswal, 2015a) to examine the whole brain characterizations of modulatory interactions effects. Another limitation of the current study is that the resting-state run was always acquired at the beginning of the scan session. We designed the tasks in this way to prevent contaminations of other tasks on the resting-state, given ample evidences that task executions can alter brain signals in resting-state (Sarabi et al., 2018; Tung et al., 2013). The order effect may contribute to the observed differences in the three conditions. Further studies may add a post task resting-state run to rule out the order effects.

In conclusion, the current analysis extended our previous analysis in resting-state and showed that the modulatory interaction among ACC and right fronto-parietal regions were highly modulated by task demands. The results may provide a new model on how error related signals affecting working memory process through higher order interactions among brain regions.

## Acknowledgements

This study was supported by grants from National Natural Science Foundation of China (NSFC61871420) and (US) National Institute of Health (R01 AT009829; R01 DA038895).

## Author contributions

X.D. conceived the idea. H.Z. designed the experiment and collected the fMRI data. X.D. performed the data analysis and wrote the draft. All authors discussed the results, and contributed to the final manuscript.

## Conflict of interest statement

The authors declare that there is no conflict of interest regarding the publication of this article.

## Data Availability

The data that support the findings of this study are available on request from the corresponding author. The data are not publicly available due to privacy or ethical restrictions.

